# CN-model: A dynamic model for the coupled carbon and nitrogen cycles in terrestrial ecosystems

**DOI:** 10.1101/2024.04.25.591063

**Authors:** Benjamin D. Stocker, I. Colin Prentice

## Abstract

Reducing uncertainty in carbon cycle projections relies on reliable representations of interactions between the carbon and nutrient cycles. Here, we build on a set of principles and hypotheses, funded in established theoretical understanding and supported by empirical evidence, to formulate and implement a dynamic model of carbon-nitrogen cycle (C-N) interactions in terrestrial ecosystems. The model combines a representation of photosynthetic acclimation to the atmospheric environment with implications for canopy and leaf N, a representation of functional balance for modelling C allocation and growth different tissues, and a representation of plant N uptake based on a first principles-informed, simplified description of solute transport in the soil. Here, we provide a comprehensive description of the model and its implementation and demonstrate the stability and functionality of the model for simulating seasonal variations of ecosystem C and N fluxes and pools. This may provide a way forward for representing key ecosystem processes related to C and N cycling based on established theoretical concepts with strong empirical support.

## 1. Introduction

Land ecosystems currently absorb around 30% of anthropogenic CO_2_ emissions and thereby slow their accumulation in the atmosphere and the rate of global heating (Friedlingstein et al., 2023). Interactions between the carbon and the nitrogen cycle (C-N interactions) in terrestrial ecosystems have the potential to limit this land C uptake under rising CO_2_ and climate change (Meyerholt et al., 2020; Thomas et al., 2013, 2015; Thornton et al., 2007; Wieder et al., 2015; Zaehle et al., 2010). N is required for the synthesis of biomass and for enzymes responsible for photosynthetic CO_2_ fixation. It has been suggested that unless additional N is available for plant uptake, additional C sequestration may be inhibited (Hungate et al., 2003).

However, C-N interactions are dynamic and plants exhibit a range of plastic responses to altered ecosystem N inputs and CO_2_ (Ainsworth and Long, 2005; De Kauwe et al., 2014; Liang et al., 2020; Medlyn et al., 2015; Pastore et al., 2019; Poorter et al., 2012). These responses have direct consequences for the degree to which N limits terrestrial C sequestration in a changing environment. Thus, general patterns in plant responses to altered C and N cycling have the potential to affect terrestrial C balance trends (Meyerholt et al., 2020). Resolving processes and capturing such general patterns in models of the terrestrial biosphere is therefore critical for reliable Earth system projections.

First principles of ecological relations have been argued to be powerful for formulating models of plant physiological processes, functional traits, growth, interactions with soil resources, and their response to changing environmental conditions (Franklin et al., 2020; Harrison et al., 2021). Eco-evolutionary optimality (EEO) principles have been successfully used for modelling and understanding different patterns and responses of ecosystem function and structure that are directly relevant for C-N interactions. These include predictions of foliar N (Dewar, 1996; Dong et al., 2022; Franklin, 2007), the share of foliar in allocated for different compartments of the photosynthetic machinery (Ali et al., 2016; Thum et al., 2019) - measured by *V*_cmax_ and *J*_max_ in the widely used Farquhar-von Caemmerer-Berry photosynthesis model (Farquhar et al., 1980), the share of C allocation to growth in roots versus shoots (Franklin, 2007; Franklin et al., 2012; Mäkelä et al., 2008; Rastetter et al., 1997; Weng et al., 2019), the investment into leaf N in the context of the water-carbon coupling (McMurtrie et al., 2008), investments into different N acquisition pathways and biological N fixation (Fisher et al., 2010; Menge et al., 2009; Rastetter et al., 2001; Wang et al., 2007), and root-rhizosphere interactions (Franklin et al., 2014; Lu and Hedin, 2019; Thurner et al., 2024).

Here, we build on the rich body of EEO-guided theoretical work for modelling C-N interactions in terrestrial ecosystems. We introduce a dynamic model that reflects a set of EEO principles and hypotheses for modelling C-N interactions - the CN-model. In contrast to some of the models underlying above-mentioned published theoretical work, the model presented here is similar in structure and scope to widely used DGVMs and components in Earth System Models that simulate land biogeochemical cycling. We combine leaf-level optimal acclimation theory (Prentice et al., 2014; Wang et al., 2017) with a set of principles that have been considered before for resolving C-N interactions in terrestrial ecosystems. The model is formulated and implemented with the aim of realistically simulating both daily C and N fluxes, comparable to ecosystem flux measurements, C and N stocks in ecosystems, and the ecosystem N balance for any terrestrial ecosystem.

We start by outlining the set of hypotheses that were considered for formulating the model presented here (Sec. 2). Then, a comprehensive model description is provided (Sec. 3) and key aspects of their implementation for dynamic simulations are described (Sec. 4). Model outputs for an example simulation are presented in Sec. 5 for demonstrating model stability and functionality.

## 2. Model hypotheses and scope

CN-model simulates the coupled ecosystem C and N dynamics in terrestrial ecosystems. It embodies a set of hypotheses that are motivated by widely observed empirical patterns and informed by eco-evolutionary optimality theory and a set of ecological and physical principles. These are are as follows.

- Carbon and nitrogen mass in the model ecosystem are conserved.
- Allocation to roots and shoots is predicted following a functional balance approach (Bloom et al., 1985) through which the root:shoot ratio is dynamically simulated such that the ratio of C assimilation and N uptake matches the demand by respiration and the C:N ratio of new biomass production.
- Acclimation of photosynthesis is modelled through the trade-off between optimising C assimilation relative to water loss (Prentice et al., 2014) and predicts *V*_cmax_ and *J*_max_, and - by implication - the total amount of leaf metabolic N per unit leaf area (Dong et al., 2022). *V*_cmax_ and *J*_max_ are assumed to be independent of soil inorganic N availability (Liang et al., 2020; Pastore et al., 2019).
- Total leaf N is the sum of metabolic and structural leaf N and each component is modelled separately (Thum et al., 2019). Canopy-total metabolic leaf N scales with acclimated *V*_cmax_ at a standard temperature of 25^*°*^C (*V*_cmax25_) and with photosynthetically active radiation absorbed by the canopy (Dewar, 1996). It thus saturates with an increasing leaf area index (LAI) of the canopy. Structural leaf N at the leaf level is modelled as a linear function of metabolic leaf N at the leaf level.
- Canopy total C assimilation scales with absorbed photosynthetically active radiation and the light use efficiency, predicted from the least-cost hypothesis and the coordination hypothesis (Prentice et al., 2014; Stocker et al., 2020; Wang et al., 2017).
- Root N uptake is governed by the inorganic soil N pool and the root biomass and is modelled as a saturating function of both, following a simplified representation of solute transport in the soil and root uptake from first principles based on McMurtrie and Näsholm (2018).
- Net N mineralisation during litter decomposition is modelled based on Manzoni et al. (2008), assuming a carbon use efficiency of (not explicitly modelled) decomposers that is treated as an empirical function of the litter C:N ratio.
- C allocated to new growth is drawn from a non-structural C (NSC) pool. Assimilated C supplies the NSC pool (Richardson et al., 2013). This buffers seasonal variations of C assimilation and partially decouples C assimilation and growth at short time scales (Cabon et al., 2022).
- C reserves are filled when the NSC pool exceeds its target and drawn from when the NSC pool gets depleted (Thum et al., 2019).
- Ecosystem N losses, subsuming gaseous and leaching losses, scale with with inorganic N pool size and with a soil temperature and a soil moisture-dependent factor.
- N fixation is not considered (in this model version).
- No tissue-level adjustment of C:N stoichimetry in wood and roots, or root N uptake capacity, and no C “overflow respiration” (Heinemeyer et al., 2012) is modelled here for re-balancing C and N acquisition and use by the plant.

## 3. Model description

### 3.1 GPP

Gross primary production (GPP) is simulated in the form of a single-big leaf light use efficiency model, corresponding to the P-model (Stocker et al., 2020), but with the leaf area index being dynamically predicted, instead of prescribed from remotely sensed observations as done in Stocker et al. (2020). The dynamics of LAI (*L* in the equations below) is governed by the leaf mass per area (Sec. 3.2.3) and C allocation to leaves (Sec. 3.4). GPP is

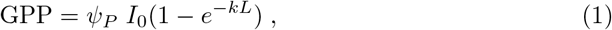

where *ψ*_*P*_ is the light use efficiency, *I*_0_ is the photosynthetically active photon flux density at the top of the canopy, *k* is the light extinction coefficient in the canopy. *ψ*_*P*_ is predicted by assuming the coordination hypothesis, which states that under typical daytime conditions, photosynthesis operates at the point of co-limitation by the capacity of Rubisco for carboxylation of RuBP and the electron transport for RuBP regeneration. Their functional forms are given by the Farquhar-von Caemmerer-Berry (FvCB) model for C_3_ photosynthesis (Farquhar et al., 1980; von Caemmerer and Farquhar, 1981). Following the electron transport-limited case, *ψ*_*P*_ can thus be written as as:

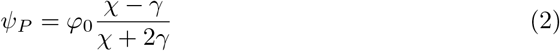

with *χ* = *c*_*i*_*/c*_*a*_ and *γ* = Γ^*\**^*/c*_*a*_, where *φ*_0_ is the quantum yield, *c*_*i*_ and *c*_*a*_ are the leaf-internal and ambient CO_2_ concentrations, and Γ^*\**^ is the CO_2_ compensation point in the absence of dark respiration, calculated after Bernacchi et al. (2001).

*χ* is predicted by accounting for an optimisation of costs associated with the carboxylation and transpiration capacities following Prentice et al. (2014). The optimal *χ* is derived following Wang et al. (2017) as

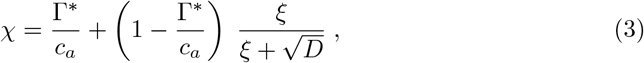

with 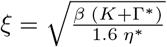 . *D* is the vapour pressure deficit and is estimated as the difference between saturated and actual vapour pressure and is a model input variable. *K* is the effective Michaelis-Menten coefficient of Rubisco. *η*^*\**^ = *η/η*_25_ where *η* is the viscosity of water and *η*_25_ is viscosity at standard conditions (25^*°*^C and 1013.25 Pa). Equation 1 can thus be written as

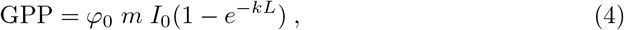

With

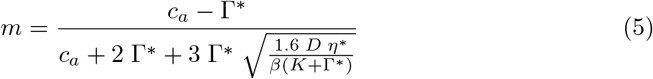

The coordination hypothesis suggests that during representative daytime conditions, the light and the Rubisco-limited assimilation rates in the FvCB model are equal. As explained in Wang et al. (2017) and Stocker et al. (2020), this can be expressed mathematically by a modification of *m* in Eq. 4:

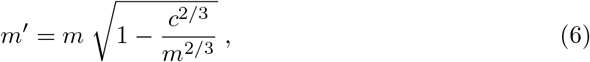

with *c* = 0.41. Finally, GPP is calculated as

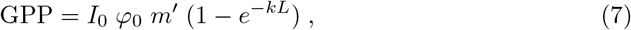

A more detailed description of the photosynthesis representation, including the temperature dependencies of Γ^*\**^, *K*, and *η*^*\**^, is provided in Stocker et al. (2020).

#### 3.1.1. V_cmax_

The maximum rate of Rubisco carboxylation, *V*_cmax_, is obtained by assuming the coordination of *V*_cmax_ and *J*_max_ such that the electron transport-limited and the Rubisco carboxylation-limited assimilation rates in the FvCB model are equal, considering average daytime conditions (see also Stocker et al. (2020), Eq. C4).

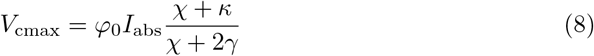

with *κ* = *K/c*_*a*_. Normalisation to the standard temperature of 25^*°*^C is done following a modified Arrhenius function based on Kattge and Knorr (2007) (see Stocker et al. (2020), Eqs. C5-C7).

#### 3.1.2. Leaf respiration

Leaf respiration is modelled as the dark respiration and is proportional to *V*_cmax25_ at standard temperature (25^*°*^C).

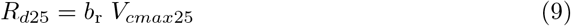

The temperature dependence of *R*_*d*_ is modelled following Heskel et al. (2016) (see also Stocker et al. (2020) Eqs. C8-C11).

#### 3.1.3. Scaling

While GPP describes the gross photosynthetic C uptake at the ecosystem, or canopy-level (i.e., per unit ground area), *V*_cmax_, and *R*_*d*_ are commonly defined at the leaf level. However, since the model is formulated as a single big-leaf model and *I*_abs_ is the light absorbed by the *canopy*, these quantities are to be understood as representative for the canopy-level. Since *V*_cmax_ and *R*_*d*_ linearly scale with *I*_abs_, and since *I*_abs_ is a saturating function of *L, V*_cmax_ and *R*_*d*_ (and canopy-total metabolic leaf N, see Sec. 3.2) also saturate with *L*.

### 3.2. Canopy N and C

N in foliage is modelled as the sum of a metabolic fraction (*N*_*v*_) and a structural fraction (*N*_*s*_).

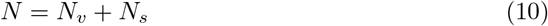

Analogously to *V*_cmax_ and *R*_*d*_, *N, N*_*v*_, and *N*_*s*_ are the canopy-total foliar, metabolic, and structural N in in foliage, respectively, expressed per unit ground area.

#### 3.2.1. Metabolic leaf N

*V*_cmax_ is defined here as a canopy-level effective quantity, representative for the single big leaf. Hence, *V*_cmax_ scales in proportion to the light absorbed by the canopy (*I*_abs_). Using the Beer-Lambert law for light absorption, a saturating relationship between (canopy total) *V*_cmax_ and the leaf area index (LAI) follows. Eq. 8 can thus be written as

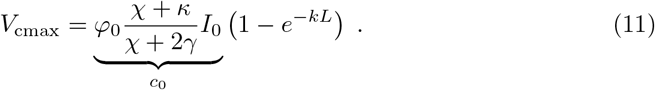

*c*_0_ is a variable substitute. The metabolic leaf N (*N*_*v*_) is taken to be proportional to *V*_cmax25_. This considers the relationship between the amount of N-rich photosynthetic enzymes, mostly Rubisco, the relationship between the amount of (active) Rubisco and *V*_cmax25_, and the coordination *V*_cmax_ and *J*_max_. The canopy-total metabolic leaf therefore also saturates with an increasing LAI.

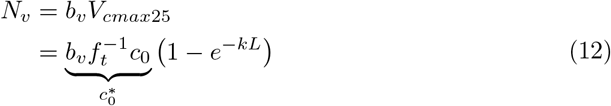

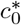 is a variable substitute. 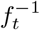 scales *V*_cmax_ to the standard temperature 25^*°*^C, following a modified Arrhenius function based on Kattge and Knorr (2007) (see Stocker et al. (2020)). At the leaf-level (denoted by the superscript *l*), the amount of metabolic N per unit leaf area is

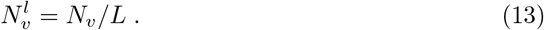

#### 3.2.2. Structural leaf N

The structural leaf N (*N*_*s*_) is modelled as a linear function of 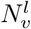.

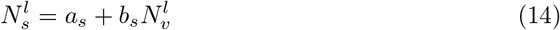

The parameter *a*_*s*_ determines the minimum investment into structures as 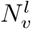 tends toward zero when *L* tends towards infinity (theoretically). The total leaf N is the sum of metabolic and structural N at the leaf level (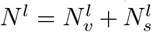). At the canopy-level, this is

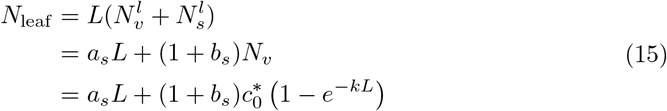

#### 3.2.3 Leaf mass per area

Leaf C per unit leaf area (*C*^*l*^) is largely equivalent to leaf mass per area, LMA, divided by two to account for the C-content in dry matter. *C*^*l*^ is calculated by multiplying *N*^*l*^ with a constant factor that relates canopy structural N to structural C.

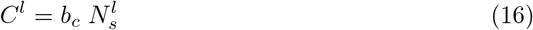

#### 3.2.4 Canopy C and N relationships

The canopy-total C in foliage, expressed as a function of LAI (*L*) is

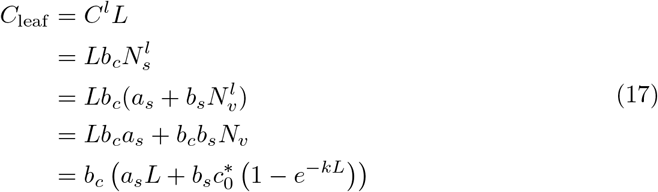

In the model, *L* needs to be calculated as a function of *C*_leaf_, e.g., after allocation. However, solving Eq. 17 for *L* is not straight-forward as the equation describes a transcendental function of the form

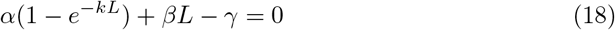

with 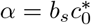, *β* = *a*_*s*_, and *γ* = *C*_leaf_ */b*_*c*_. To express *L* as a function of *C*_leaf_, the Lambert-W function *W*_*n*_ has to be used:

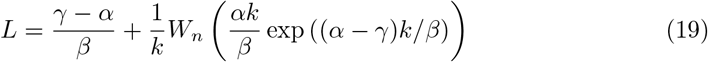

Taken together, these relationships imply a non-linear relationship between *C*_leaf_ and LAI (Fig. 1). With an increasing LAI, the C:N ratio increases, and LMA decreases (not shown). While the total metabolic N component is proportional to *V*_cmax25_ at the canopy-level and thus reaches an asymptote with increasing LAI, the minimum amount of N (determined by *a*_*s*_) and C (determined by *b*_*c*_) needed for foliage tissue always implies a non-zero cost for adding more leaves.

**Figure 1.**
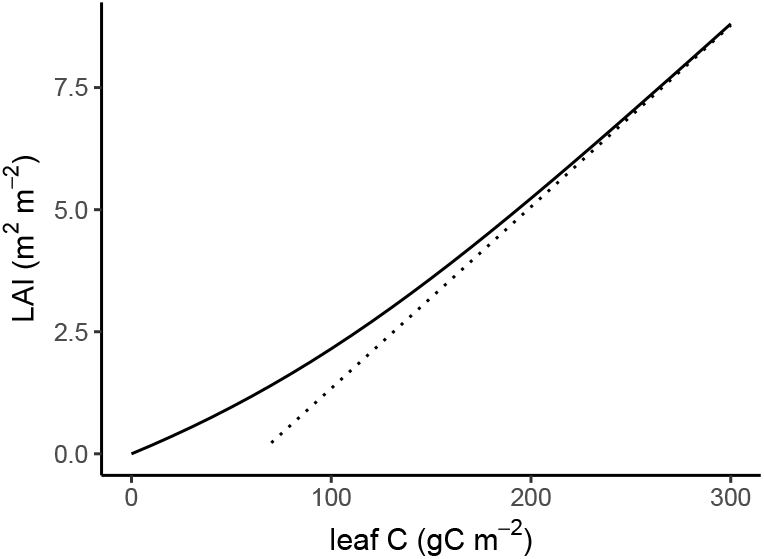
Illustration of the relationship between LAI and leaf C for an example case with *a*_*s*_ = 0.056, *b*_*s*_ = 1.23, *b*_*C*_ = 40. The dashed line is the asymptote 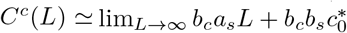.

#### 3.2.5 Simplified canopy C and N

It may be assumed that there is no relationship between the amount of metabolic and structural N in leaves. To reflect this, the parameter *b*_*s*_ can be set to zero. Thus, the non-linear relationship between *L* and *C*_leaf_ becomes linear.

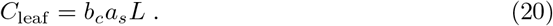

### 3.3 Non-structural C and N dynamics

Non-structural C (NSC) is modelled with two pools - a labile pool (*C*_labl_) and a reserves pool (*C*_resv_). *C*_labl_ is fuelled by net C assimilation (GPP - *R*_*d*_) and depleted by C used for root respiration (*R*_*r*_), sapwood respiration (*R*_*w*_), C exudation (*C*_ex_), C allocated to new growth (*C*_alc_) and C allocated to the reserves pool Δ*C*_resv_. Respiration terms are described in Sec. 3.3.1. *C*_labl_ can also be replenished by a flux from the reserves pool into the labile pool (Δ*C*_resv_ *<* 0, see Sec. 3.4.3). In each model time step, the change in *C*_labl_ is given by:

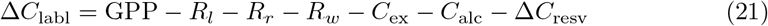

If C mass in the labile pool is insufficient to satisfy respiratory demands (*C*_labl_ *< R*_*l*_ + *R*_*w*_ + *R*_*r*_ + *C*_ex_, terms introduced in Sec. 3.3.1), the remainder is withdrawn from the reserves pool *C*_resv_. Exuded C is added to a separate exudates pool from where it is respired without further effects (Sec. 3.7).

Non-structural N (NSN) is modelled analogously to NSC, but with plant N uptake (*N*_up_) as the source of N and without withdrawals from respiration, and assuming that no N is exuded.

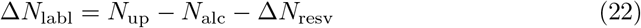

*N*_up_ is described in Sec. 3.5. *N*_alc_ is described in Sec. 3.4.1. N is moved between the labile and the reserves pool (Δ*N*_resv_) in parallel to C, described in Sec. 3.4.3.

#### 3.3.1 Autotrophic respiration and exudation

Leaf respiration is taken to be equal to canopy-total dark respiration (Eq. 9).

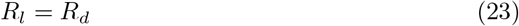

Root maintenance respiration is modelled as being proportional to root C mass and an air temperature-dependent factor.

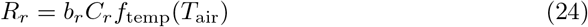

Sapwood respiration is modelled analogously.

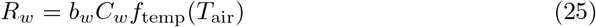

The temperature-dependent factor is a modified Arrhenius equation (Lloyd and Taylor, 1994), adopted from the LPJ model (Sitch et al., 2003). Its functional form deviates from an exponential representation as represented via the common Q10 coefficient for chemical reactions and implicitly accounts a temperature acclimation of respiration.

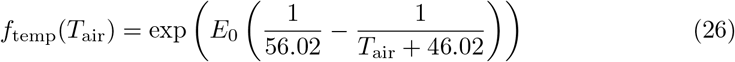

*E*_0_ is set to 308.56 J (Sitch et al., 2003).

Exudation of labile C by roots into the rhizosphere is treated by a separate temperature-independent C flux.

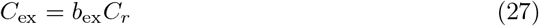

### 3.4 Allocation

#### 3.4.1 C and N for new growth

The amount of C allocatable to new growth is proportional to the size of the labile C pool (*C*_labl_) and is constrained by the amount of N available in the labile N pool (*N*_labl_).

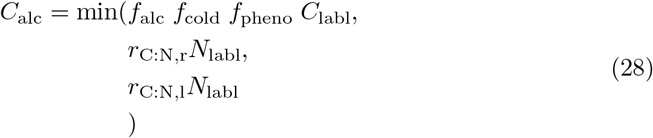

Due to the generally higher C:N ratio in woody biomass than in leaves and roots, its value (*r*_C:N,w_) does not constrain the allocation more than it does when considering *r*_C:N,r_ and *r*_C:N,l_, and is therefore not considered in Eq. 28. *f*_alc_ is set to 0.1. *f*_cold_ is a unitless factor that varies between zero and one and is reduced by freezing temperatures, and increases as a function of temperature sums (growing degree days). This introduces a lag in the resumption of vegetation growth to rising temperatures after cold days, but an instantaneous suppression of growth by low temperatures. It is implemented as described in Appendix A1 and in the Supplementary Note S1 in Luo et al. (2023). *f*_pheno_ is set to 1.0 and thus takes no effect in this model version. *r*_C:N,r_ is the C:N ratio in root biomass and is set as a constant (model parameter). *r*_C:N,l_ is the C:N ratio in leaf biomass and is determined by the formulation described in Sec. 3.2. N is allocated to new growth in parallel to C, considering the constant root and wood C:N ratio (*r*_C:N,r_, *r*_C:N,w_) and the dynamic leaf C:N ratio (Sec. 3.2.4).

#### 3.4.2 Functional balance

Allocation to roots and shoots is modelled following a C-N functional balance approach through which the fraction of C allocated to leaves *f*_*l*_ is dynamically simulated such that the ratio of C assimilation and N uptake (left side of Eq. 29) matches the demand by respiration and the C:N ratio of new biomass production (right side of Eq. 29).

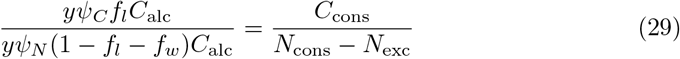

*y* is the growth efficiency (after accounting for growth respiration) and cancels out in Eq. 29 (as does *C*_alc_). (1 − *f*_*l*_ − *f*_*w*_) is the fraction allocated to roots (after accounting for the fractions allocated to leaves and wood, *f*_*l*_ and *f*_*w*_, respectively). *ψ*_*C*_ is the efficiency of net C assimilation (GPP - *R*_*d*_) per unit C allocated to leaves. *ψ*_*N*_ is the efficiency of plant N uptake per unit C allocated to roots. Eq. 29 thus implicitly builds on linearized relationships, diagnosed from allocation and acquisitions over the previous 365 days (*d*).

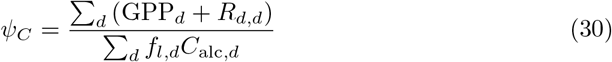

The efficiency term *ψ*_*N*_ is calculated as the ratio of running sums of N uptake and and the amount of C allocated to root growth, summed over the preceding 365 days.

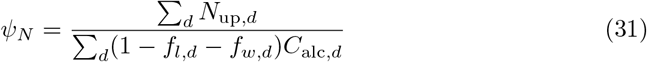

*C*_cons_ comprises C allocated to new growth, to growth respiration (accounted for my multiplication with 1*/y*), and to root and sapwood respiration (*R*_*r*_ and *R*_*w*_), and C exudation (*C*_ex_), calculated as the running sums over the preceding 365 days (*d*).

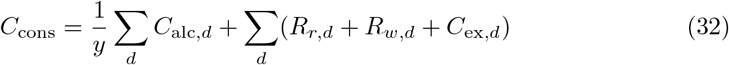

*N*_cons_ comprises N allocated to new growth, summed over the preceding 365 days.

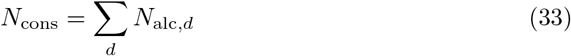

*N*_exc_ accounts for the difference between total acquired N (by root uptake) and the N consumed for new growth, summed over the preceding 365 days.

The allocation algorithm uses Eq. 29, solved for *f*_*l*_, to determine the fraction of allocatable C (*C*_alc_), invested to new leaf growth:

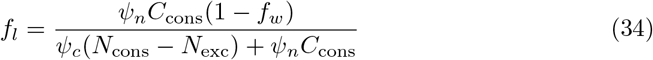

The fraction of allocatable C invested into wood growth is taken to be a constant *f*_*w*_. The remainder (1 − *f*_*l*_ − *f*_*w*_) gets allocated to root growth.

#### 3.4.3 Allocation to reserves

Δ*C*_resv_ in Eq. 21 is modelled similarly as in Thum et al. (2019) as the net of a “pull” into the reserves pool (*f*_lr_) and a “push” in the opposite direction (*f*_rl_).

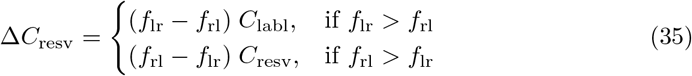

The “pull”, *f*_lr_, is a sigmoid function of the ratio of the current reserves pool over its target size 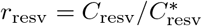.

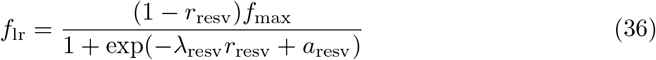

*a*_resv_ is set to -1.5. *f*_max_ is set to 0.02 d^−1^. The target size of the reserves pool 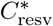 is proportional to the total amount of C allocated to new growth over the preceding 365 days (*d*).

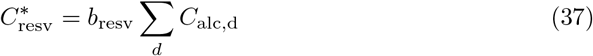

The “push”, *f*_rl_ scales linearly with *r*_resv_ and non-linearly with the ratio of the current labile pool over its target size 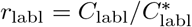.

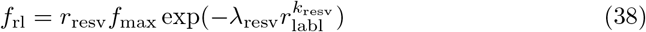

*k*_resv_ is set to 3.0 and *λ*_resv_ to 10.0.

The target size of the labile pool 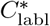 is proportional to the total amount of C consumed by respiration and exudation over the preceding 365 days (*d*).

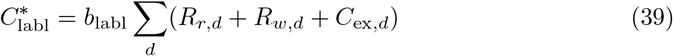

*k*_resv_, *λ*_resv_ and *a*_resv_, and therefore shape of the push and the pull functions, are chosen such that the the flux between the labile and the reserves pool (Δ*C*_resv_) is zero most of the time. It is non-zero and positive only if the labile pool is substantially larger than its target size, while the the reserves pool is smaller than its target size. It is negative non-zero only if the labile pool is close to being depleted while C is available in the reserves pool. These dynamics are illustrated in Fig. 2.

**Figure 2.**
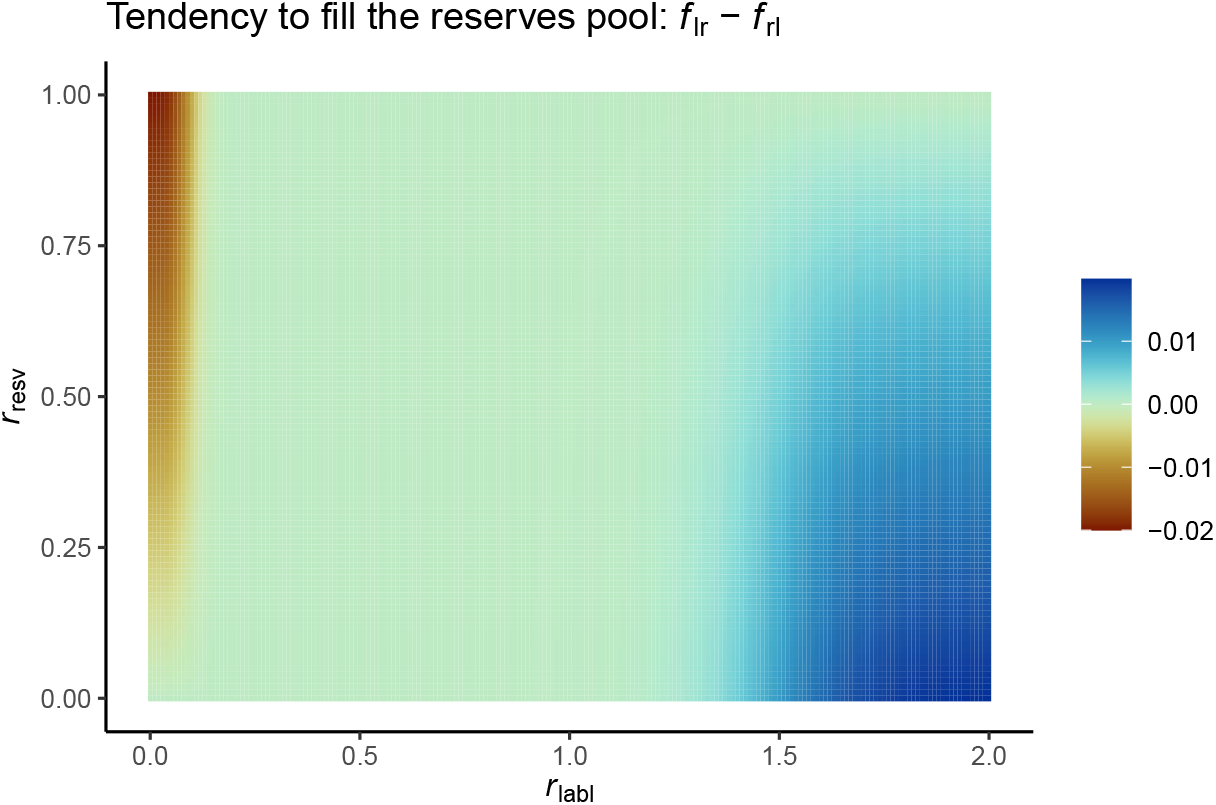
Tendency to fill the reserves pool (*f*_lr_ − *f*_rl_) as a function of the ratio of the labile pool size over its target size (*r*_labl_), and of the reserves pool size and its target size (*r*_resv_).

**Figure 3.**
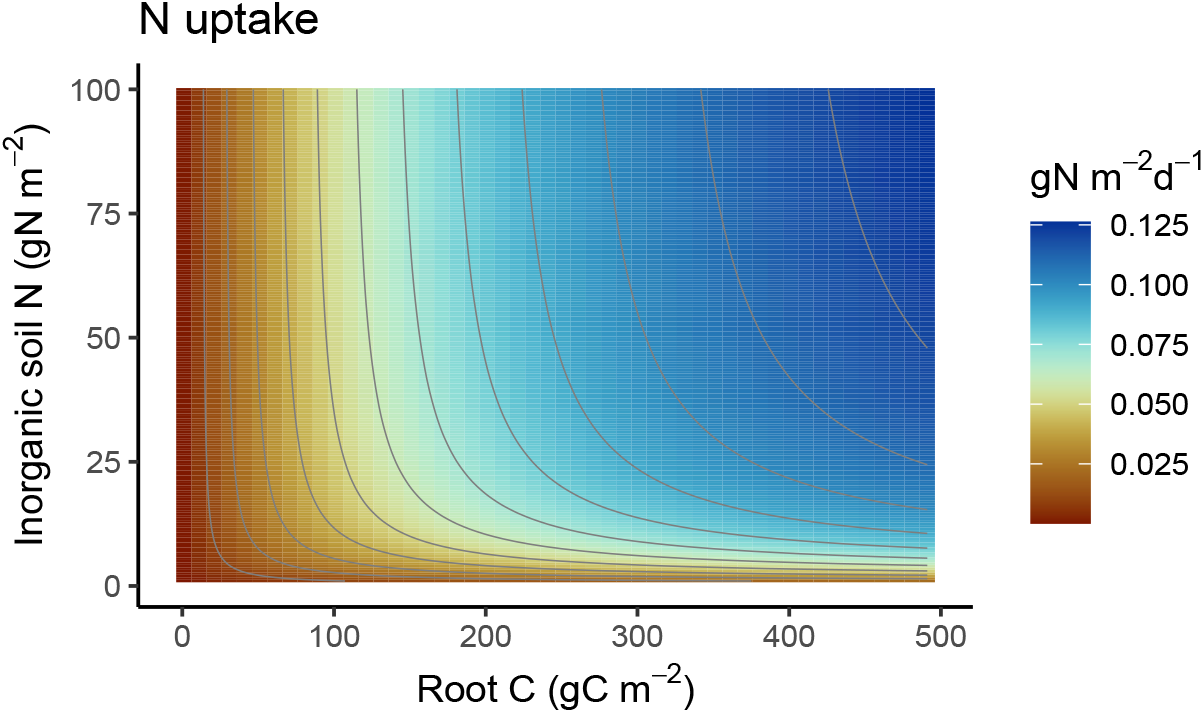
Nitrogen uptake as a function of root C and soil inorganic N. The model is run with parameters *V*_max_ = 5 gN m^−2^d^−1^, *k*_*C*_ = 250 gC m^−2^, and *k*_*V*_ = 5 gN m^−2^.

Nitrogen gets transferred between the labile and reserves pools along with carbon, carrying the C:N ratio of the respective pool from which C is drawn.

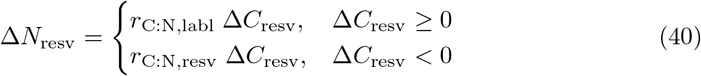

### 3.5 Nitrogen uptake

McMurtrie and Näsholm (2018) developed a model for solute transport and root N uptake via diffusion and advection for representation in terrestrial biosphere models. Their model resolves the link between root biomass, morphological traits, and the capacity for nutrient uptake, subject to root water uptake. Their model also implies that N uptake saturates (for a given set of root morphological traits) both with respect to root biomass and inorganic N in the soil solution. The CN-model implements a simplification of the model by (McMurtrie and Näsholm, 2018) and inherits its general functional response to soil inorganic N and root biomass based on Michaelis-Menten kinetics. The CN-model does not account for the link between water and nutrient flows and does not explicitly resolve the role of mass-specific root surface area but retains the key characteristic of the model by (McMurtrie and Näsholm, 2018) of a simultaneous saturation of plant N uptake with respect to root biomass (*C*_root_) and soil inorganic N (*N*_inorg_). Nitrogen uptake is modelled as the product of two saturating functions:

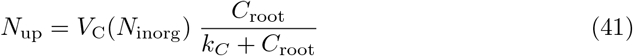

and

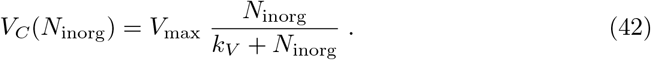

*V*_max_ is an uptake capacity at saturating soil inorganic N. *V*_*C*_(*N*_inorg_) is an *N*_inorg_-dependent uptake capacity at saturating root mass. Root N uptake dynamics are described in McMurtrie and Näsholm (2018) per unit soil volume. Here, we do not resolve the vertical soil profile, but consider the whole soil column as a homogenous well-mixed volume and express all pools and fluxes per unit ground area.

### 3.6 Biomass turnover

Biomass turnover follows first-order kinetics (Olson, 1963). The fraction of biomass turned over at each model time step (day) is modelled as

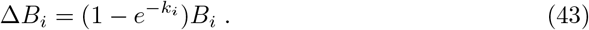

Note that for *k*_*i*_ *≪* 1, 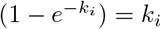, but the form of Eq. 43 is chosen to avoid numerical instability for cases where *k ≪* 1 does not apply. Since C and N are turned over in parallel, we describe dynamics here for *B*_*i*_ = (*C*_*i*_, *N*_*i*_) and all equations are valid for both the respective C and N pools. *i* refers to the leaf, root, wood, seed, or labile pool. *k*_*i*_ is its respective decay rate constant. An exception from this is that a fraction *f*_resorb_ of the leaf turnover is retained (resorbed from senesced leaves) and stored in the labile N pool.

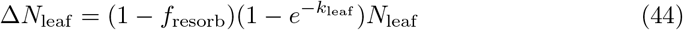

At each model time step, biomass pools get updated as

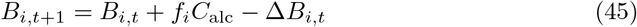

*f*_*i*_*C*_alc_ is the biomass production (Sec. 3.4.1). The turned over biomass Δ*B*_*i,t*_ is added to the litter pool. Woody biomass turnover is added to an aboveground litter pool with slow turnover rates (‘lags’), leaf turnover are added to an aboveground litter pool with fast turnover rates (‘lagf’), root turnover is added to a belowground litter pool (‘lbg’).

### 3.7 Soil organic matter dynamics and net N mineralisation

#### 3.7.1 Decomposition

This model part is largely adopted from LPJ (Sitch et al., 2003) and DyN-LPJ (Xu-Ri et al., 2008), but the C use efficiency of the litter-to-SOM transfer is modified as a function of the litter C:N ratio after Manzoni et al. (2008). For completeness, we describe the full set of equations used in the CN-model. Analogously as above, we refer to *P*_*i*_ = (*C*_*i*_, *N*_*i*_) (gC m^−2^ and gN m^−2^) as the pools, containing both C and N, and the decomposition of litter and soil pools turn over C and N in parallel. Pools are the three litter pools described above and two soil pools (fast ‘sf’, slow ‘ss’).

Soil organic matter (SOM) and litter turnover are modelled as first-order decay processes with decay rate constants modified by soil moisture (*θ*) and soil (air) temperature (*T*) for SOM and belowground (aboveground) litter decomposition.

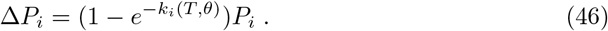

The decay constants *k*_*i*_ (d^−1^) are calculated as the product of a base respiration rate *k*_0,*I*_ (Tab. 1), modified by an empirical function *f*_*θ*_ of soil moisture (*θ*) after Foley (1995) and a modified Arrhenius function *f*_*T*_ of soil temperature *T*_soil_ after Lloyd and Taylor (1994)

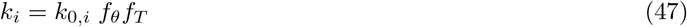

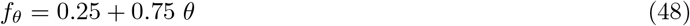

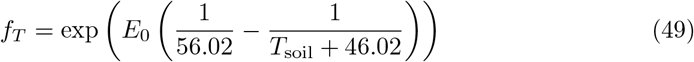

The equation for *f*_*T*_ is identical to the one used for modelling the temperature dependency of tissue maintenance respiration (Eq. 26).

At each model time step, the litter and soil pools get updated as

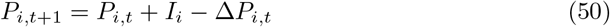

*I*_*i*_ is the input flux of C and N into pool *P*_*i*_. For litter pools, this is the biomass turnover:

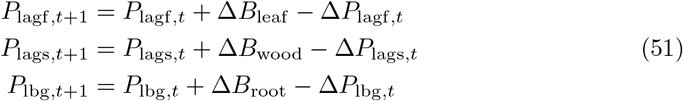

For the litter-to-SOM transfer, C and N dynamics are modelled separately (see below).

#### 3.7.2 C dynamics

The fates of decomposed litter C are determined by the C use efficiency of the litter-to-SOM transfer (*e*_C_) and the diversion fractions (*f*_sf_, 1 − *f*_sf_) of decomposing litter to two SOM pools (*P*_sl_, *P*_sf_) with different turnover times (*k*_sl_, *k*_sf_):

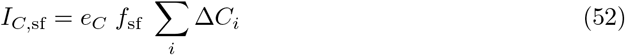

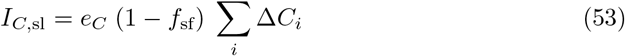

where *i* are three litter pools *i* = (lfs, lsl, lbg) The remainder of decomposing litter is respired as CO_2_ and is counted towards heterotrophic respiration. The C use efficiency *e*_*C*_ is calculated here based on an empirical relationship derived from litter bag studies (Manzoni et al., 2008) between the N:C ratio of decomposing litter (*r*_*L*_) and the critical litter N:C ratio (*r*_CR_) above which net mineralisation occurs (net immobilisation prevails below *r*_CR_):

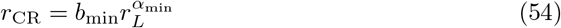

This implies that decomposers with a N:C ratio of *r*_*B*_ adjust their C use efficiency *e*_*C*_ so that

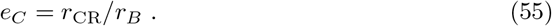

*r*_*L*_ is dynamically simulated subject to the C:N ratio of turned-over biomass, N resorption efficiencies (see Sec. 3.6), and relative shares of individual biomass compartments to litter formation. *r*_*B*_ represents the decomposers’ N:C ratio and is a taken as a constant here (see Tab. 1).

**Table 1.** Model parameters. ‘Code name’ refers to the parameter name in the model code. Values are chosen either by adoption from the the P-model Stocker et al. (2020), the LPX-Bern model Stocker et al. (2013), or chosen here such that simulated pools and fluxes are broadly consistent with observations.

| Parameter | Code name | Value | Reference |
| --- | --- | --- | --- |
| **GPP** |  |  |  |
| $\varphi_0$ | kphio | 0.08 | Eq. 2 |
| $k$ | kbeer | 0.5 | Eq. 1 |
| **Canopy N and C** |  |  |  |
| $b_v$ | nv_vcmax25 | 273.63 | Eq. 12 |
| $a_s$ | ncw_min | 0.056 | Eq. 14 |
| $b_s$ | r_n_cw_v | 1.23223 | Eq. 14 |
| $b_c$ | r_ctostructn_leaf | 40 | Eq. 16 |
| **NSC and NSN** |  |  |  |
| $b_r$ | r_root | 1.826 | Eq. 24 |
| $b_r$ | r_sapw | 0.088 | Eq. 25 |
| $b_{ex}$ | exurate | 0.003 | Eq. 27 |
| **Allocation** |  |  |  |
| $f_{alc}$ | frac_avl_labl | 0.1 | Eq. 28 |
| $r_{C:N,r}$ | r_cton_root | 37.0 | Eq. 28 |
| $r_{C:N,w}$ | r_cton_wood | 350 | |
| $y$ | growtheff | 0.6 | Eq. 29 |
| $f_w$ | frac_wood | 0.5 | Eq. 29 |
| **Nitrogen uptake** |  |  |  |
| $V_{max}$ | nuptake_vmax | 1.0 | Eq. 42 |
| $k_C$ | nuptake_kc | 250 | Eq. 41 |
| $k_V$ | nuptake_kv | 1.0 | Eq. 42 |
| **Biomass turnover** |  |  |  |
| $k_{leaf}$ | k_decay_leaf | 1 | Eq. 43 |
| $k_{root}$ | k_decay_root | 1 | Eq. 43 |
| $k_{wood}$ | k_decay_sapw | 0.01 | Eq. 43 |
| $k_{labl}$ | k_decay_labl | 0 | Eq. 43 |
| $f_{resorb}$ | f_nretain | 0.5 | Eq. 44 |
| **SOM dynamics** |  |  |  |
| $k_{0,lags}$ | klitt_af10 | 1.2 yr<sup>-1</sup> | Eq. 49 |
| $k_{0,lagf}$ | klitt_as10 | 0.35 yr<sup>-1</sup> | Eq. 49 |
| $k_{0,lbg}$ | klitt_bg10 | 0.35 yr<sup>-1</sup> | Eq. 49 |
| $k_{0,sf}$ | ksoil_fs10 | 0.021 yr<sup>-1</sup> | Eq. 49 |
| $k_{0,ss}$ | ksoil_sl10 | 0.0007 yr<sup>-1</sup> | Eq. 49 |
| $f_{sf}$ | | 0.985 | Eq. 60 |
| $b_{min}$ | ntoc_crit1 | 0.45 | Eq. 54 |
| $\alpha_{min}$ | ntoc_crit2 | 0.76 | Eq. 54 |
| $1/r_B$ | cton_soil | 9.77 gC (gN)<sup>-1</sup> | Eq. 60 |
| **Inorganic N dynamics** |  |  |  |
| $k_{0,N}$ | maxnitr | 0.00005 | Eq. 63 |

#### 3.7.3 Net N mineralisation

Litter gross N mineralisation is determined by the N content in decomposing litter. This can be written as

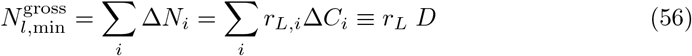

with *i* = (laf, las, lbg). The litter net N mineralisation is determined by the gross mineralisation rate and the critical N:C ratio below which net mineralisation is negative (net immobilisiation):

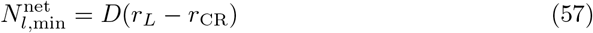

In the model, litter decomposition generally implies N immobilisation as *r*_CR_ *> r*_*L*_ in most cases. N required for immobilisation depletes the soil inorganic N pool. N is transferred to the SOM pools at a N:C ratio of *r*_*B*_. This implies that decomposers impose their N:C ratio immediately upon all newly formed SOM. N input into the fast and slow SOM pools is hence:

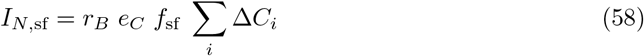

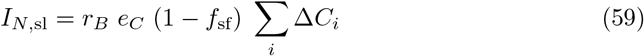

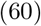

with *i* = (*laf, las, lbg*). This further implies that SOM gross N mineralisation doesn’t cause any further immobilisation of N as SOM has the same N:C ratio as decomposers and all N mineralised from SOM decomposition is transferred to the inorganic soil N pool and thus becomes available for plant N uptake.

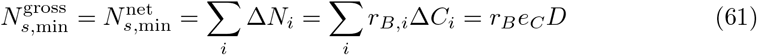

with *i* = (ss, sf). Note that decomposers are not represented by an explicit pool here. The last equality holds in equilibrium where SOM C inputs are equal to decomposing C that is not respired (*e*_*C*_*D* = Δ*C*_*i*_ for *i* = (ss, sf)). From this it follows that in steady state, the total (litter plus soil) net N mineralisation is 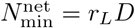.

### 3.8 Inorganic N dynamics

Inorganic N dynamics (Δ*N*_inorg_) are determined by the balance of net N mineralization from litter (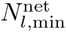, Eq. 57) and soil (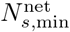, Eq. 61) decomposition, losses through gaseous and leaching pathways, and inputs by atmospheric deposition or other (e.g., N fertilisation in ecosystem experiments, *N*_dep_).

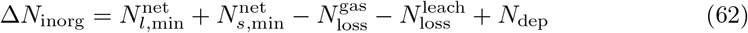

Both loss pathways scale in proportion with the soil mineral N pool size. Gaseous losses are modelled as a function of soil temperature after Lloyd and Taylor (1994) (*f*_*T*_, Eq. 49):

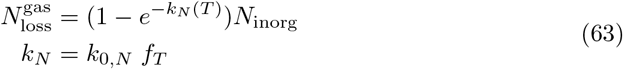

Leaching losses are modelled to scale with the ratio of daily soil water runoff to total root zone water storage. The soil water balance is modelled following Davis et al. (2017). *N*_dep_ is a model forcing.

## 4 Implementation

The model is dynamic, operates at daily time steps, tracks all C and N fluxes and transfers between pools, and conserves C and N mass. It is forced by time series of daily values for meteorological conditions (temperature, precipitation, vapour pressure deficit, photosynthetically active radiation, CO_2_, and N-deposition or other forms of reactive N inputs to the soil).

### 4.1 Functional balance under seasonal variations

C fixation by photosynthesis and N uptake from the soil varies greatly over the course of a year in most terrestrial ecosystems. Yet, the growth of plant organs responsible for resource acquisition (e.g., leaves for C fixation, fine roots for N uptake) is slow. Thus, the size of plant organs responsible for the different resource acquisitions may not be attuned to a perfect functional balance under rapidly varying environmental conditions. This poses a challenge for modelling C-N interactions.

Two measures are implemented here to model a “damped” response of the ecosystem to rapid variations in the environment. First, the maximum *V*_cmax25_ over the preceding 365 days is considered for determining metabolic leaf N and 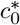 in Eq. 12. Second, C and N acquisition of the preceding 365 days is considered for determining the functional balance of allocation to leaves versus roots as described in Sec. 3.4.2.

### 4.2 Code

The model is implemented as an R package in the {rsofun} modelling framework and relies on low-level source code, implemented in a set of Fortran 90 modules, containing subroutines for modelling individual processes (Fig. 5). All modelled quantities are contained in Fortran 90 *derived types* (a list of variables of different types and dimensions). Quantities with memory (C, N pools, and water pools) and quantities without memory between time steps (fluxes) are contained in separate derived types. Organic pools (biomass, litter, and SOM) are defined as *derived types*, containing C (and its isotopes, currently only ^12^C) and N (and its isotopes, currently only ^14^N). Functions for transferring organic mass (biomass, litter, SOM) simultaneously transfer their different constituents.

**Figure 4.**
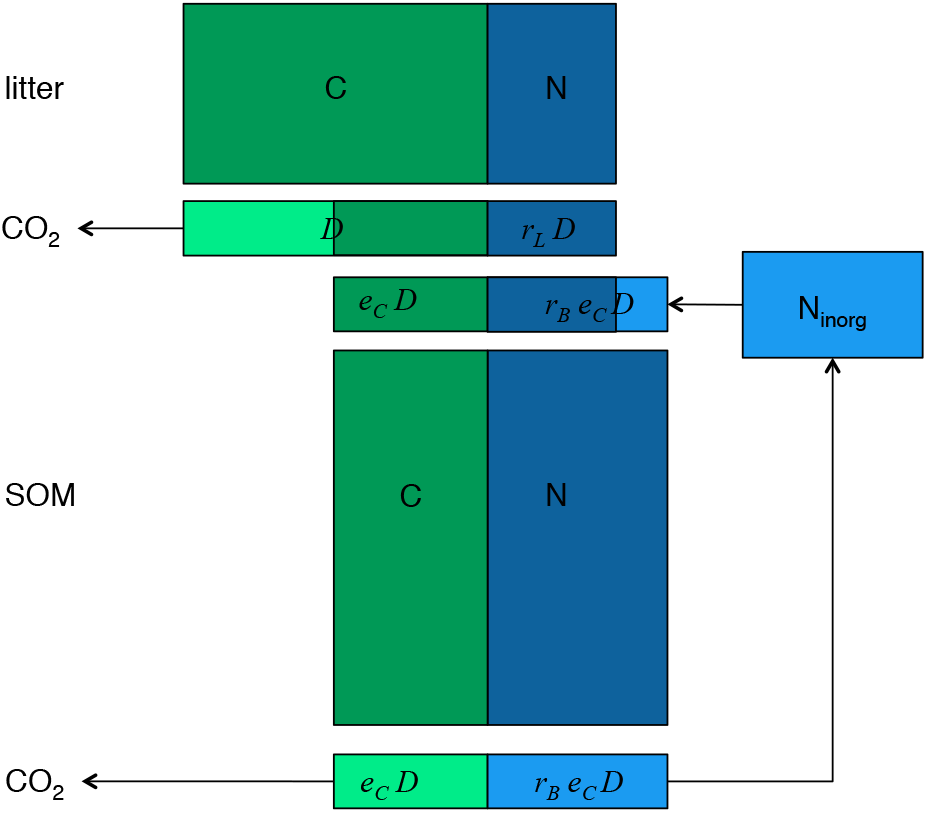
Litter and soil decomposition and nitrogen mineralisation. *D* is the decom-posing litter C mass at each time step. *e*_*C*_ is the carbon use efficiency of decomposers and the soil organic matter pool (SOM). *r*_*L*_ is the N:C ratio of litter. *r*_*B*_ is the N:C ratio of decomposers and SOM. Light green indicates C that gets respired as CO_2_. Light blue indicates mineralised N. Net mineralisation is the mineralisaiton of SOM deccomposition minus the immobilisation by litter decomposition.

**Figure 5.**
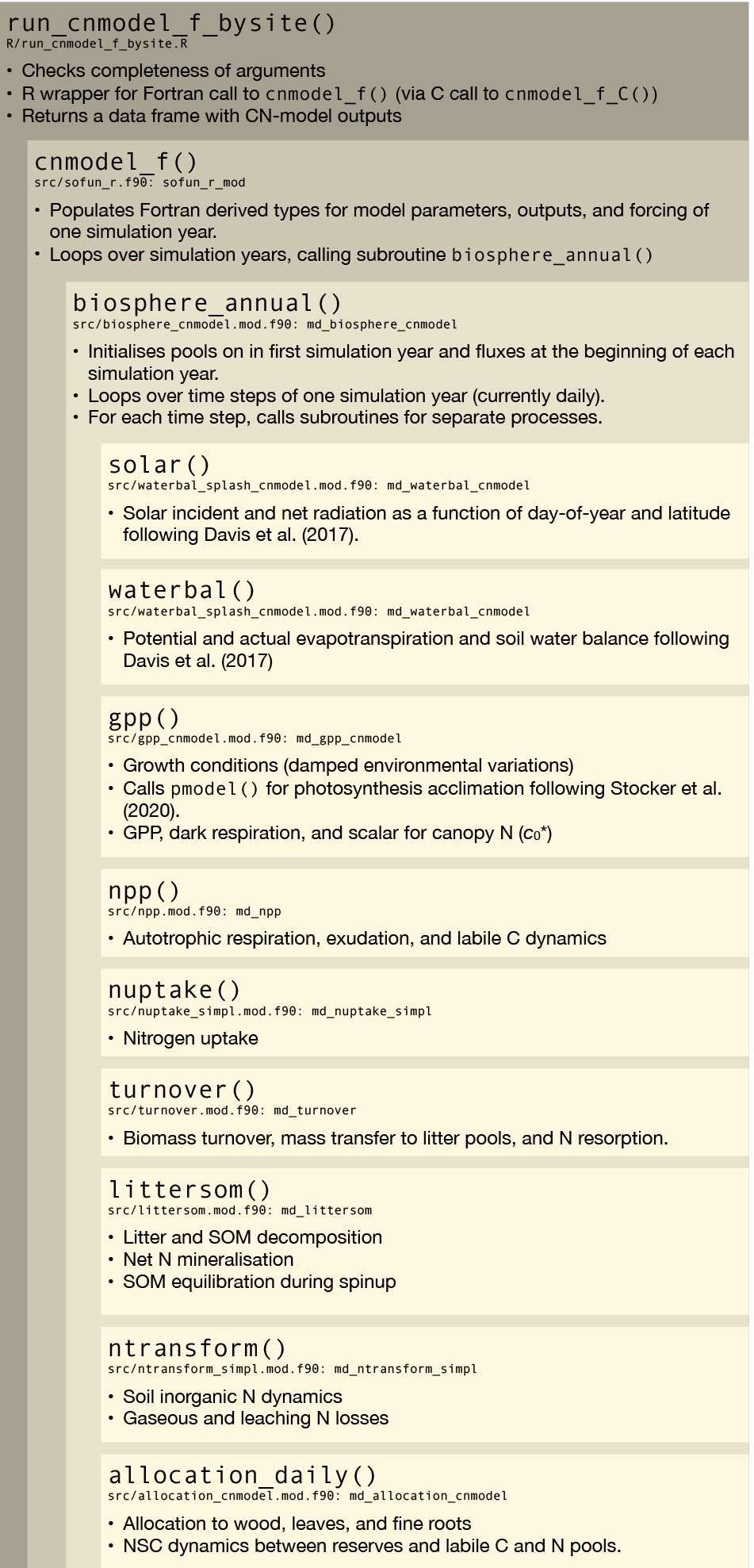
Model architecture describing the nested structure of major functions and subroutines implemented in different source code files.

### 4.3 Spin-up

Soil organic C pools are spun up to a steady state at a constant forcing (repeating one year of model forcing). Soil organic C pools are set to an analytically determined steady-state twice during the model spin-up. Steady-state SOC pool sizes 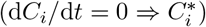 are estimated based on averaged input and decomposition rates (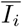 and 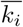, respectively) during the preceding five years:

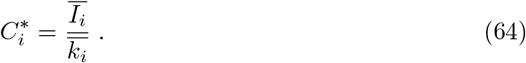

Soil organic N pools are then set based on 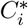 and the C:N ratio of the pools during the preceding five years before the analytical soil C equilibration was applied.

Due to the feedback between ecosystem productivity, biomass turnover, soil organic carbon, net N mineralisation, soil inorganic N availability, and (again) ecosystem productivity, special model spin-up procedures have to be imposed and successively relieved to attain a steady state in an accelerated manner. The procedures are as follows and their sequence during the model spin-up is illustrated in Fig. 6.

**Figure 6.**
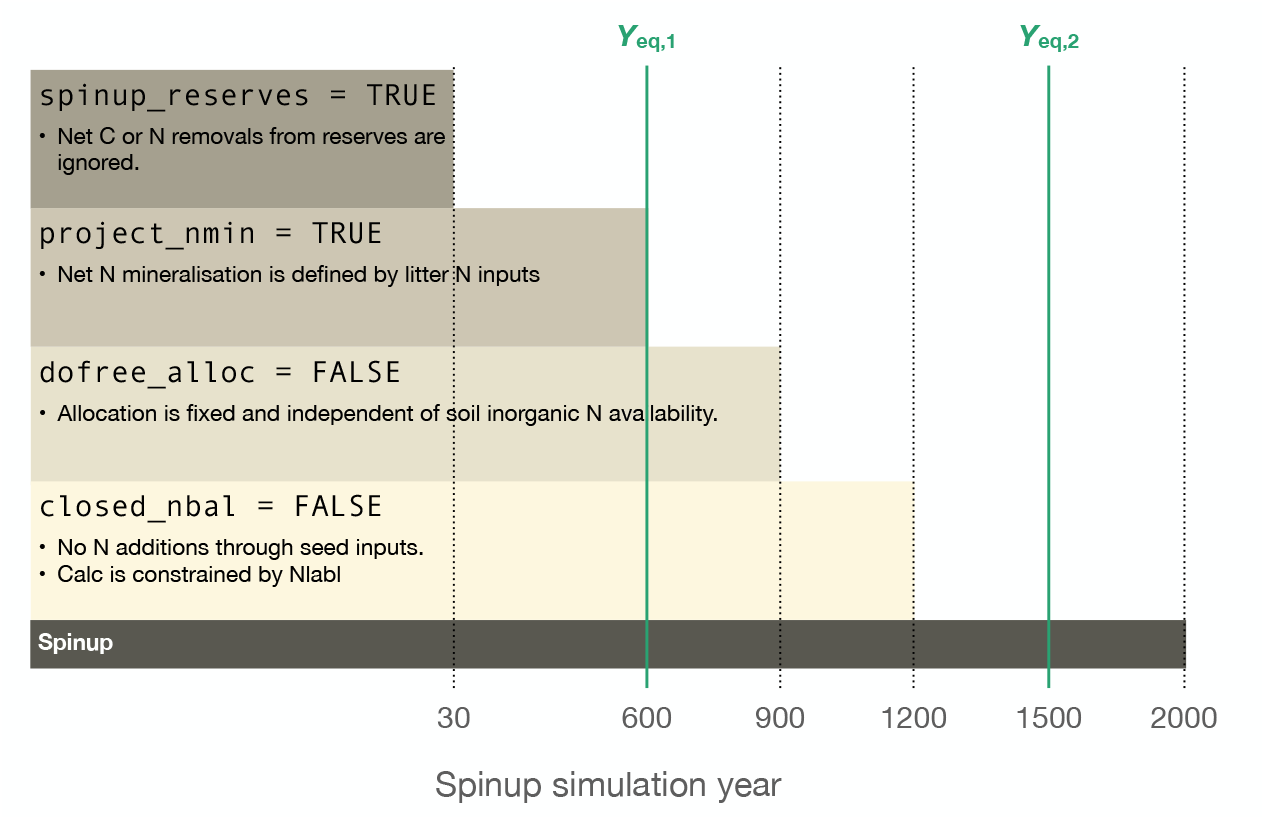
Model spin-up levels. The end of the different spin-up procedures are indicated by the dotted lines. Timings (*Y*_eq,1_ and *Y*_eq,2_) of the two analytical SOM equilibrations are indicated by the green lines.

#### Fill reserves (spinup reserves = TRUE)

Net C or N removals from reserves are ignored, altering Eq. 35 to:

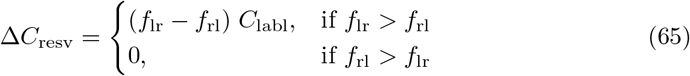

#### Projecting net N mineralisation (project nmin = TRUE)

Net N mineralisation is modelled as a function of N inputs into the litter pools. This is done by assuming that in steady-state, N inputs to litter must equal net N mineralisation from litter and SOM decomposition:

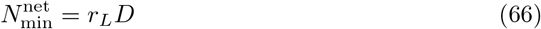

This avoids the “vicious circle” of critically low net N mineralisation rates before new N accumulates in the SOM pool that would inhibit vegetation growth and thereby prevent SOM accumulation.

#### Fixed allocation (dofree alloc = FALSE)

Allocation is fixed (set as model parameter (`frac leaf`) and independent of soil inorganic N availability.

#### Ignoring closed N balance (closed nbal = FALSE)

N (and C) is added as seed biomass (addition to labile pools) as specified by the model forcing. Runtime model checks to conserve the N balance and avoid negative values for N pool sizes are ignored. Allocation of N to new growth may “over-deplete” the non-structural N pool, thereby effectively fixing N and violating N mass conservation.

### 4.4 Simulation protocol

The model is forced with a mean seasonal cycle of meteorological variables, derived from data obtained from a temperate grassland ecosystem flux tower site in Switzerland (CH-Oe1). Meteorological model forcing variables include air temperature, vapour pressure deficit, photosynthetic photon flux density (PPFD), atmospheric pressure, precipitation, atmospheric CO_2_, and cloud cover fraction. Annual atmospheric N deposition is obtained from **?** and evenly divided into daily portions (amounting to annual totals of 1.5 gN m^−2^ yr^−1^). Atmospheric CO_2_ is held constant at 380 ppm. The model is spun up for 2001 years and run for seven years (years 2002-2008 in output).

Model parameter values are set as described in Tab. 1. In particular, the wood allocation fraction is set to zero

## 5 Results

### 5.1 Spin-up

The spin-up of the soil organic C pools is shown in Fig. 7. The first analytical pool equilibration in spin-up simulation year 600 (left vertical green line in Fig. 7) brings pools to near their magnitude attained at the end of the spin-up. Soil C pools increase slightly after the relief of the fixed allocation phase in spin-up year 900 and decline again slightly after relief of the un-closed N balance phase in spin-up year 1200 (dotted lines in Fig. 7). The second analytical soil C pool equilibration is in spin-up year 1500 (right vertical green line in Fig. 7). As shown in Fig. 7b, soil C pools are not at a perfect steady-state, even after the second analytical equilibration. This is due to the feedback between soil C pool size, net N mineralisation and availability, NPP, biomass turnover, and (again) soil C pool size. However, the remaining trend in the soil C pools at the end of the spin-up (last 100 years of model spin-up) is small and amounts to -0.0002% yr^−1^.

**Figure 7.**
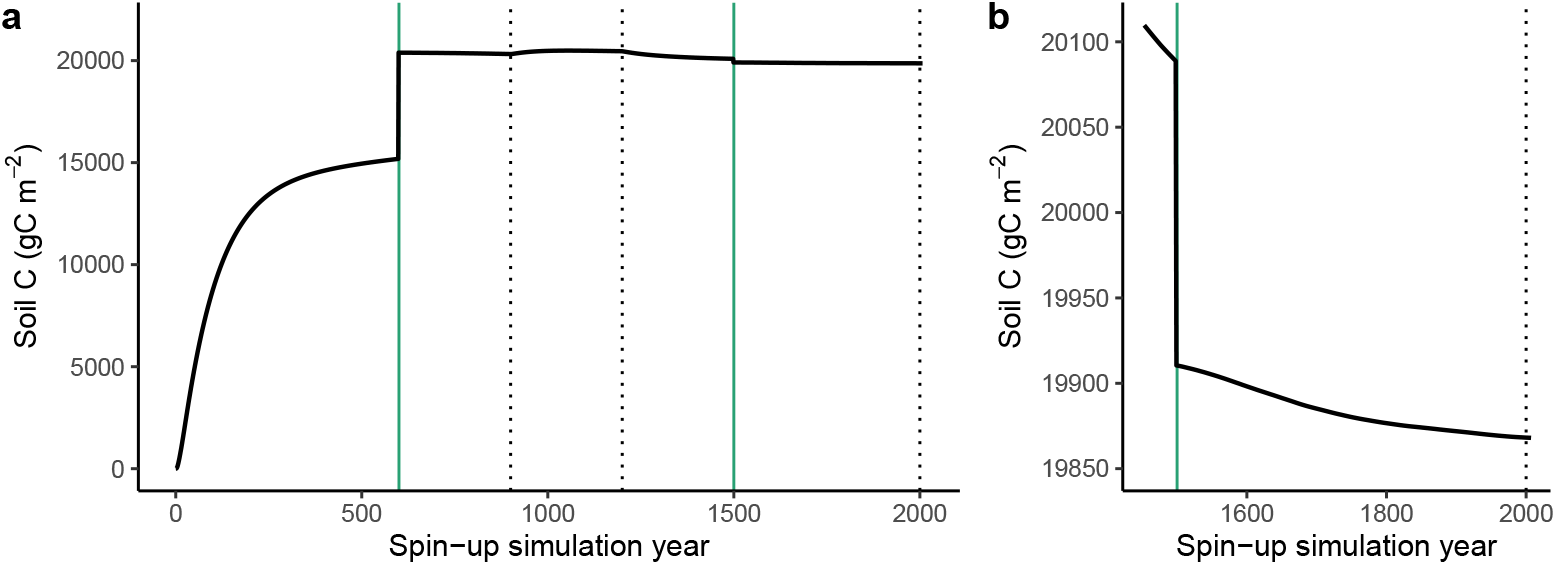
Simulated soil C during the entire model spin-up (a) and a zoomed-in section of spin-up years 1450-2000 (b). Vertical lines correspond to vertical lines in Fig. 6. Vertical green lines indicate the timings of the analytical soil C equilibration. Vertical dotted lines indicate timings when spin-up procedures are relieved (first vertical dotted line: fixed allocation, second vertical dotted line: un-closed N balance) and the end of the model spin-up (third vertical dotted line).

### 5.2 Time series

Time series for multiple simulated variables over seven model simulation years after the spin-up is shown in Figs. 8 and 9. In response to seasonal variations in meteorological conditions, most variables exhibit a clear seasonal pattern. For each variable, the same seasonal pattern is repeated each year, reflecting that the model forcing is a repeated mean seasonal cycle and that the model state is dynamic, yet stable.

**Figure 8.**
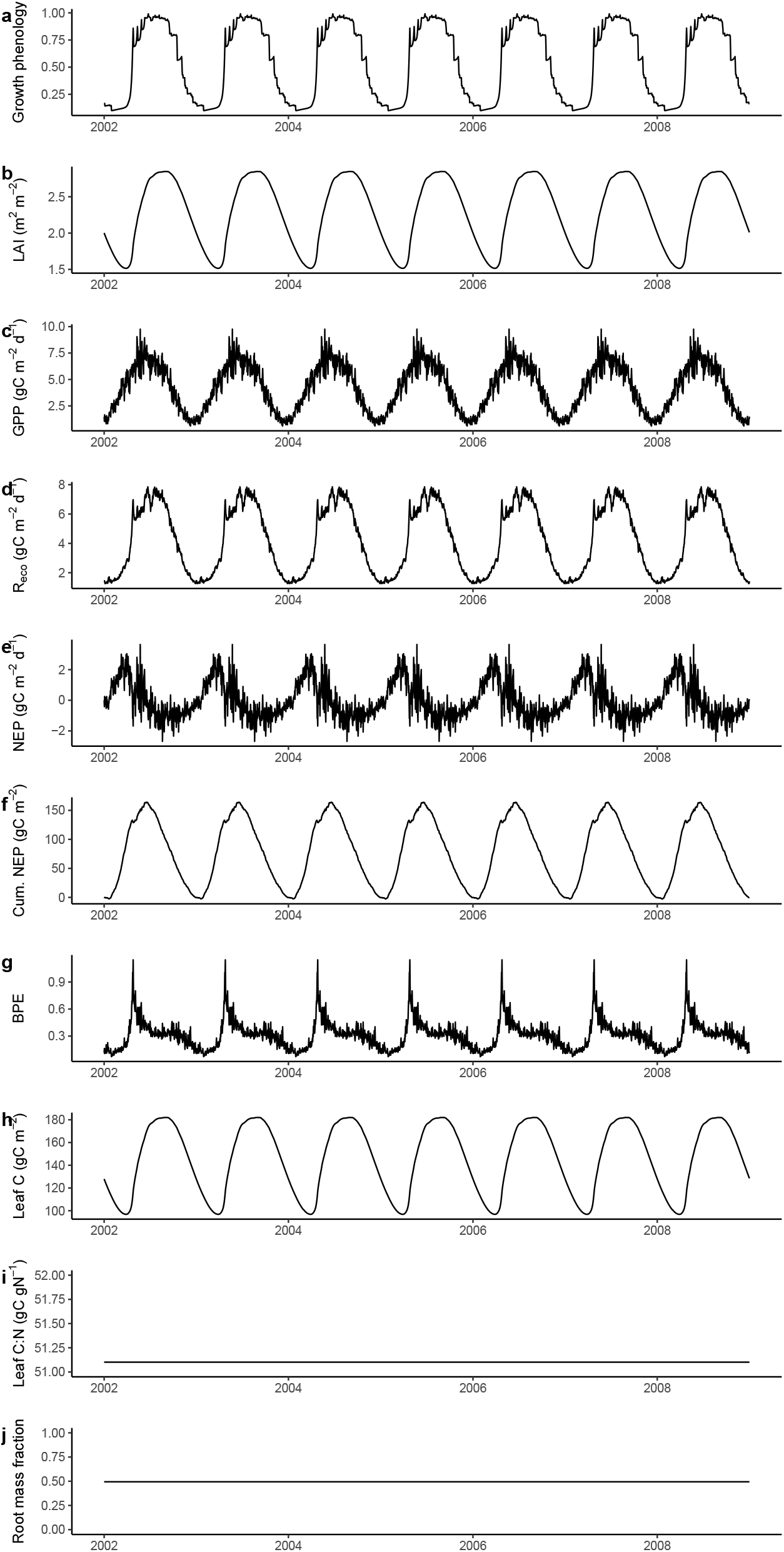
Simulated time series of (a) vegetation growth phenology, (b) leaf area index, (c) gross primary production, (d) ecosystem respiration (*R*_eco_), (e) net ecosystem productivity (NEP), (f) the cumulative sum of NEP, (g) biomass production efficiency calculated as (Δ*C*_leaf_ + Δ*C*_root_ + Δ*C*_wood_ + Δ*C*_seed_)*/*GPP, (h) leaf C, (i) leaf C:N, and (j) the root mass fraction.

**Figure 9.**
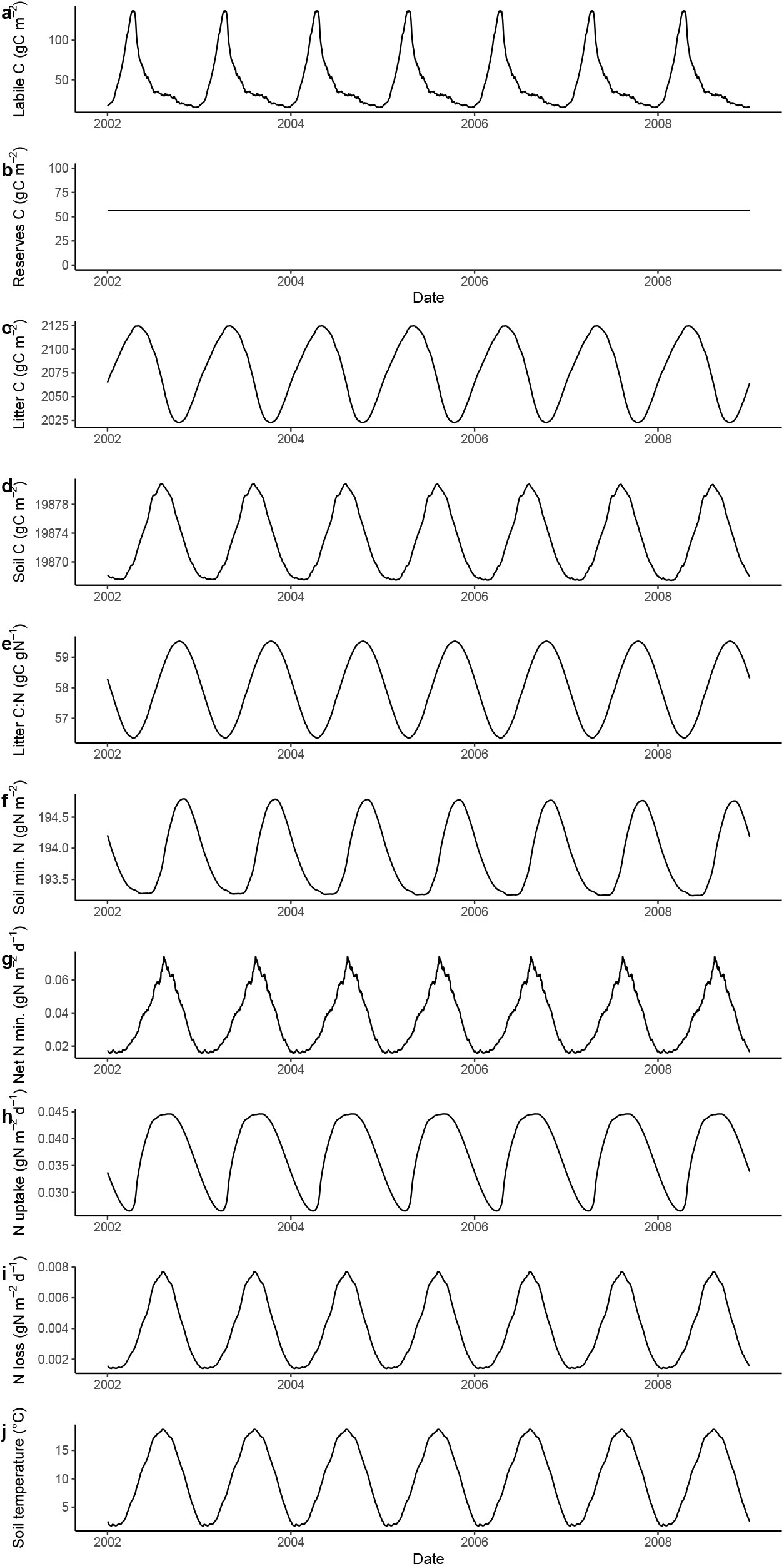
Simulated time series of (a) C in the labile pool, (b) C in the reserves pool, (c) litter C, (d) soil C, (e) litter C:N, (f) soil inorganic (mineral) N, (g) net N mineralisation, (h) plant N uptake, (i) total ecosystem N losses as the sum of gaseous and leaching losses, and (j) soil temperature.

Vegetation growth phenology (Fig. 8a) is reduced during winter and increases rapidly in spring, triggering the rapid development of leaf area, as expressed by the seasonal variation in the leaf area index (Fig. 8b). The seasonal pattern of GPP (Fig. 8c) is determined by the patterns in LAI and PPFD (not shown). Net ecosystem productivity (NEP, Fig. 8e), computed from model outputs as GPP − *R*_*a*_ − *Rh*, peaks earlier than GPP due to the delayed peak of ecosystem respiration *R*_eco_ = *R*_*a*_ + *Ra* (Fig. 8d) compared to the seasonal peak of GPP. The cumulative sum of daily NEP values (Fig. 8f) has a seasonal pattern, but no visually discernible long-term trend. This reflects the steady-state of the ecosystem C balance and the complete accounting of the ecosystem mass balance by the fluxes in the model output (GPP, *R*_*a*_, *R*_*h*_).

The ratio of biomass production (BP = Δ*C*_leaf_ + Δ*C*_root_ + Δ*C*_wood_ + Δ*C*_seed_) to GPP is referred to as the biomass production efficiency (BPE). It varies strongly within a year (Fig. 8g). This reflects the fact that biomass production (allocation of C to growth) is governed not only by the balance of GPP and *R*_*a*_ supplying *C*_labl_, but also by the seasonally varying growth phenology.

Leaf C (Fig. 8h) and N vary in parallel with LAI, such that the leaf C:N ratio is stable (Fig. 9d). Also the leaf C and root C (not shown) vary in parallel, such that the root mass fraction is stable (Fig. 8j). The labile C pool peaks in spring - before the onset of the vegetation growth phenology and the increase in LAI which draws down C in the labile pool (Fig. 9a).

In contrast to most variables, some variables exhibit no seasonal pattern. Besides the leaf C:N and the root mass fraction, also the reserves pool is stable (Fig. 9b), reflecting that a net flux from and to the reserves pool occurs only under extreme conditions (Fig. 2). Since the model is forced with the same mean seasonal cycle in each simulation year, no such extreme condition occurs. The root mass fraction is stable because the balance of C and N acquisition, accumulated over 365 days is considered for modelling allocation (Sec. 3.4.2), thereby evening out sub-annual variations in C and N acquisition. Similarly for the leaf C:N ratio. It is modelled as a function of the metabolic leaf N content which scales with (seasonally varying) *V*_cmax25_. However, the *maximum V*_cmax25_ over the preceding 365 days is considered for determining metabolic leaf N (via determining 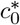 in Eq. 12).

The litter C:N ratio exhibits a seasonal variation due to the input of fresh litter (with a relatively high C:N ratio) during the growing season and the progressive respiration of litter C and immobilisation of mineral N during litter decay (Fig. 4). Seasonal variations in soil mineral N (Fig. 8f) reflect the balance of N uptake (Fig. 8h), net N mineralisation (Fig. 8g), and N losses (Fig. 8i). The latter two are controlled by soil temperature (Fig. 8j).

### 5.3 Allocation and mass balance

The mean seasonal cycles of the vegetation C and N balances are shown in Figs. 10 and 11. C is acquired by vegetation through photosynthesis (GPP) and consumed by biomass production, respiration, and exudation, and lost by a turnover of the labile pool. The C consumption and loss is referred to as C ‘sinks’ in Fig. 10b. Considering annual totals, C sinks equal GPP. This follows from mass balance conservation and the aspect that the simulated ecosystem is in a dynamic equilibrium with no long-term (multi-annual) accumulation of C in any of the vegetation pools. Yet, within a year, GPP and the sinks are not in perfect alignment (Fig. 10a). The mismatch of C sinks and sources at a daily time scale is balanced by plant-internal storage in the form of the labile and the reserves pool in this model. C is assimilated in excess of consumption by the sinks in the early part of the year. During the rapid leaf and root growth phase in spring, collective C consumption by all sinks exceeds GPP by a factor of *∼*2 and is slightly higher than GPP on several days during the remainder of the year. The cumulative excess C consumption exactly matches the cumulative excess C assimilation such that annual totals are balanced (Fig. 10b).

**Figure 10.**
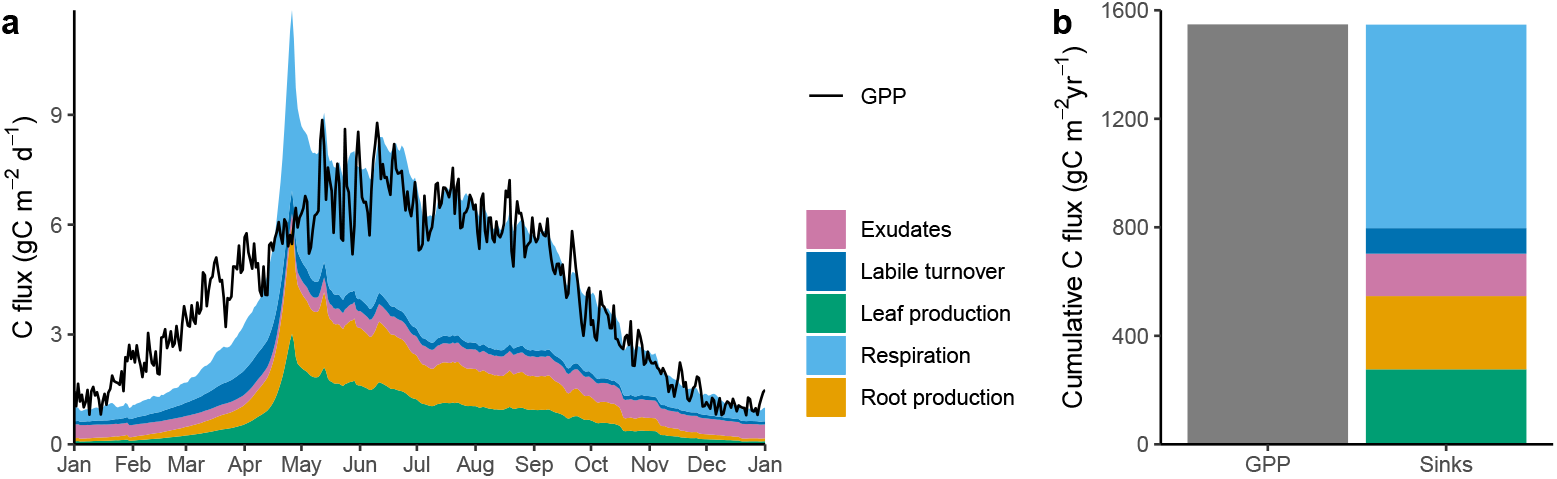
Mean seasonal cycle of the vegetation C balance. (a) Seasonal course of GPP (black line) and all processes consuming C. Colored regions are additive and their total area represents annual totals. (b) Annual totals of GPP and all processes consuming C (‘sinks’).

**Figure 11.**
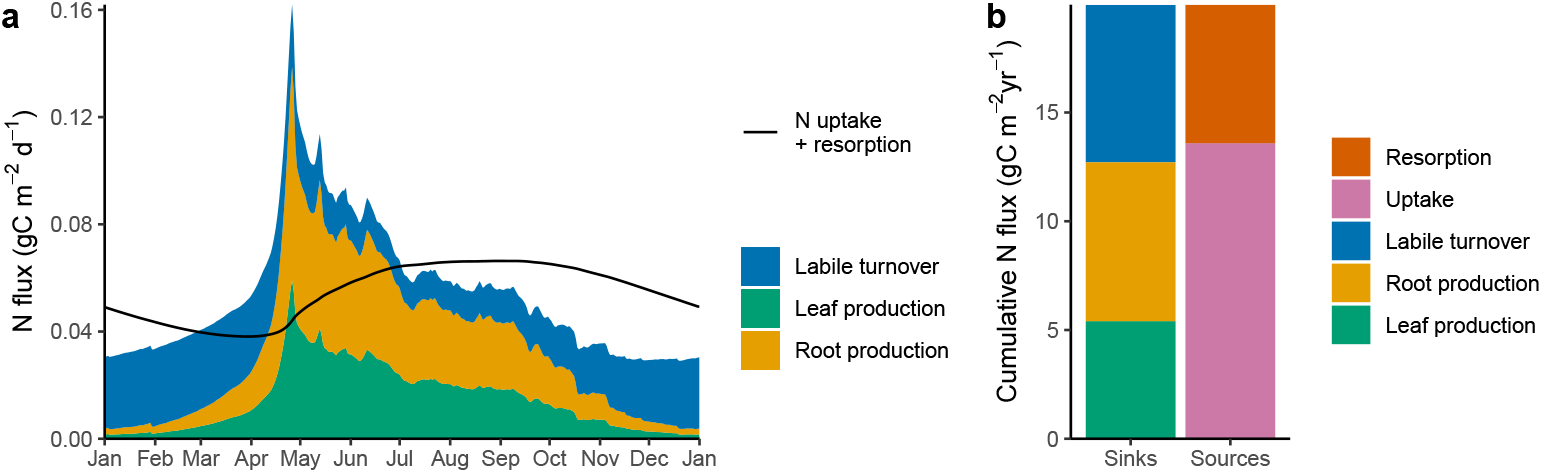
Mean seasonal cycle of the vegetation N balance. (a) Seasonal course of the sum of N uptake and resorption (black line) and all processes consuming N (biomass production). Colored regions are additive and their total area represents annual totals. (b) Annual totals of the sum of N uptake and resorption (‘sources’) and all processes consuming N (‘sinks’).

**Figure 12.**
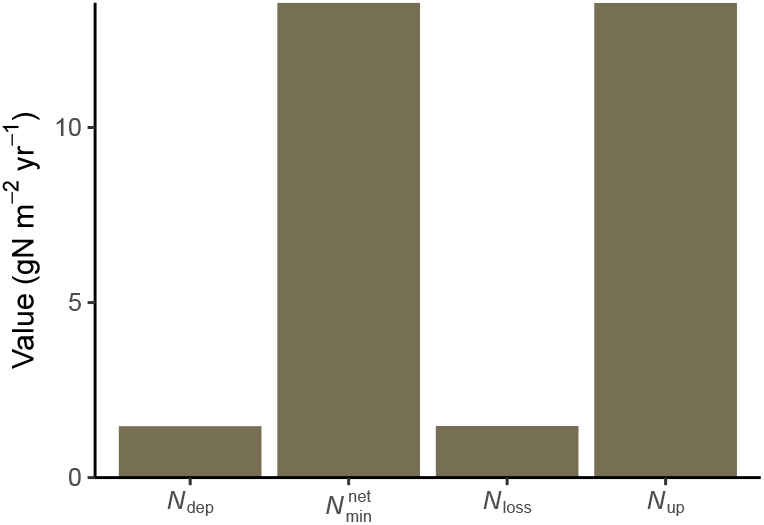
Annual ecosystem N fluxes. *N*_dep_ is atmospheric N deposition, 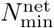 is net N mineralisation, *N*_loss_ is the total of gaseous and leaching N losses, *N*_up_ is N uptake.

The same general pattern is simulated for N. Here, ‘sinks’ are biomass production and a labile N loss term, no exudation of N is considered in this model, and respiration terms are absent. N ‘sources’ are plant uptake from the soil inorganic N pool, and resorption from senescing tissues. The mismatches between sinks and sources a the daily time scale are even larger for N than for C (Fig. 10a). The largest N sink occurs in spring at the onset of the vegetative growth phenology. Also N is balanced by plant-internal storage in the labile and reserves pool such that the annual total sinks and sources match.

### 5.4 N balance

The model simulates that, considering annual totals, most N is recycled within the ecosystem (Fig. A1). The magnitude of N losses matches the magnitude of external inputs through atmospheric deposition. The magnitude of plant N uptake matches the magnitude of net N mineralisation. This reflects the steady-state of ecosystem N cycling in the simulation. The ratio of losses (or N deposition)) to net N mineralisation (or N uptake) is a measure for N cycle openness (Cleveland et al., 2013) and is 11% here.

## 6 Discussion

We have demonstrated here how a set of eco-evolutionary optimality principles and other fundamental principles of plant function and biogeochemical cycling can be considered for formulating a dynamic model of C-N interactions in terrestrial ecosystems - similar in scope and structure to widely used Dynamic Global Vegetation models (Friedlingstein et al., 2023; Prentice et al., 2007) and the land carbon and nutrient cycle components of Earth System Models (Davies-Barnard et al., 2020). The model presented here combines the capability of resolving ecosystem fluxes, daily plant growth, and foliage development in response to seasonal variations in the environment (Fig. 8) with a flexible allocation and a balanced C and N acquisition, attuned to long-term stoichiometric constraints and growth in different plant tissues.

The stoichiometric balance between C and N acquisition and use (growth and respiration) is simulated as being governed by a flexible allocation to leaves and roots. Unlike in other models, the C:N stoichiometry of biomass is assumed to not respond to an imbalance in C and N acquisition versus. In the model presented here, plasticity in foliar C:N stoichimetry follows as a consequence of acclimation of photosynthesis to the growth environment and canopy development - without a direct influence of soil N. Furthermore, the non-structural C and N pools, modelled with a *labile* pool and a *reserves* pool, are crucial for balancing mismatches between C and N acquisition and use that arise in response to short-term (sub-annual) variations in the environment (Figs. 10 and 11). This implies a partial decoupling between photosynthesis and growth at sub-seasonal time scales (Fig. 8g). However, at longer (multi-annual) time scales, the link between photosynthesis and growth persists as there is no “overflow respiration” of C (Heinemeyer et al., 2012) or loss of N for re-balancing the C and N supply versus demand. As a consequence of the functional balance assumption for modelling allocation, root growth declines if labile N is available in excess. If labile C is available in excess, leaf growth declines.

The sensitivity of allocation and the root:shoot ratio (or the root mass fraction) to C and N (experimental treatment of CO_2_ and ecosystem N inputs) has strong empirical support (Ainsworth and Long, 2005; Cleland et al., 2019; De Kauwe et al., 2014; Eastman et al., 2021; Jiang et al., 2020; Keller et al., 2023; Leakey et al., 2009; Li et al., 2020; Poorter et al., 2012; Rogers et al., 1995; Schneider et al., 2004; Song et al., 2019). The model presented here may provide a basis for predicting these experimentally observed responses. The demonstration of such model predictions and their evaluation against observations is beyond the scope of this study. The model representation of non-structural C and N pools is motivated by findings of an important role of non-structural C in plants for sustaining critical function under stressed conditions (Richardson et al., 2013) and for supplying the resource for rapid foliage development in deciduous plants.

The assumed absence of an influence of soil inorganic N availability on tissue C:N ratios, foliar N content, and photosynthetic capacities (*V*_cmax_ and *J*_max_) constitutes a deviation from common approaches taken in C-N interactions-resolving DGVMs. It is motivated by the following observations and findings. First, the response of *V*_cmax_ to experimental N fertilisation is often found to be relatively weak or highly variable across experiments (Liang et al., 2020). This contrasts with the general pattern of clear influence of soil N on foliar N (Liang et al., 2020) and suggests that photosynthetic capacities (*V*_cmax_ and *J*_max_) are only partially coupled to foliar N content. Second, *V*_cmax_ can be predicted to decline under elevated CO_2_ merely as a consequence of photosynthetic acclimation to the atmospheric environment (Smith and Keenan, 2020). In an ecosystem experiment, this decline has been found to not be alleviated by N fertilisation (Pastore et al., 2019). Third, the role of stoichiometric flexibility has been found to be overestimated in models and this overestimation has been linked to deficiencies in their simulations of the responses of ecosystem N cycling under elevated CO_2_ (Medlyn et al., 2015; Zaehle et al., 2014). Unreliable simulations of stoichiometric flexibility have important implications for C and N cycles and may undermine projections of the land C balance and water cycling to future climate and CO_2_ changes (Hauser et al., 2023; Meyerholt et al., 2020).

The leaf and canopy-level foliar C and N relations are simulated here by resolving distinct controls on metabolic and structural components of foliar N (Harrison et al., 2009). The underlying principle is that the photosynthetic capacity for Rubisco carboxylation (*V*_cmax_) and the amount of (N-rich) Rubisco is controlled by light levels in the canopy and the total amount of photosynthetically active radiation absorbed by the canopy (Dewar, 1996). Within-canopy gradients in light levels and foliar N are not resolved by the big-leaf representation of the CN-model. The non-linear relationship between total foliar C and N simulated by the CN-model (Fig. 1) thus implicitly assumes a retranslocation of N across the canopy in response to LAI variations and leads to a simulated relationship between Rubisco-N and absorbed photosynthetically active radiation that is consistent with an optimal distribution of leaf N in the canopy (Dewar, 1996). This prediction should be empirically tested.

The modelling of interactive C and N cycling with flexible allocation and limited stoichiometric flexibility introduces additional challenges compared to a C-only model setup. The feedback between soil N availability, vegetation growth and N uptake, turnover, and N mineralisation (plant-soil feedback), and the discrepant time scales of involved processes leads to system dynamics that imply practical challenges. One such challenge is the model spin-up (Sec. 4.3). We have demonstrated here how a C-N interaction model can be spun up from empty initial C and N pools to a (near) steady-state. Another challenge is that not all model parametrisations and setups with different forcings yield a solution where C and N pools are positive and non-zero. That is, under certain boundary conditions (not shown), vegetation activity and positive C and N pools are not sustained after the relief of the spinup phases (fixed allocation, ignoring closed N balance, Fig. 6). This behaviour appears to be linked to the plant-soil feedback. It will be important to identify a parameter space that enables stable model integration under the full range of terrestrial environmental conditions where vegetation is sustained as observed.

The plant-soil feedback, combined with the non-linearity of key processes (e.g., light absorption as a function of LAI, N uptake as a function of fine root mass and soil N availability) also poses a challenge for model parameterisation. Parameters of individual processes (Tab. 1) cannot be constrained by related observations in isolation. All model parameters influence all simulated ecosystem fluxes, pools, and (due to the non-linearity of key processes) sensitivities to environmental conditions. Furthermore, each model parameter combination is associated with a distinct model steady-state and thus requires a separate (computationally costly) model spin-up. An effective calibration of the CN-model parameters and a comprehensive evaluation against a range of observations is an open task and is not addressed in this paper.

The model presented here is made freely accessible and can be used as an R package, wrapping computationally efficient low-level Fortran 90 code. The process representation in the model is designed in a modular fashion and lends itself to adoption into other modelling frameworks. In particular, the representation of flexible allocation is formulated in a manner that makes it independent of the representation of other modelled processes and does not rely on computationally costly iterations. This may provide a way forward for representing a key process based on a well-established theoretical concept (functional balance, Bloom et al. (1985)) with strong empirical support, by adopting the implementation presented here.

## 7 Conclusions

Reducing uncertainty in carbon cycle projections relies on reliable representations of interactions between the carbon and nutrient cycles. Stoichiometric flexibility has been identified to be overestimated in several C-N interaction-resolving DGVMs, while the flexibility of allocation is often underestimated or not considered in such models. We showed here how a set of principles and hypotheses, funded in established theoretical understanding and supported by empirical evidence, can be used for designing a dynamic model of C-N interactions in terrestrial ecosystems. The model presented here combines a representation of flexibility in allocation, guided by a functional balance of N and C uptake and use, with a representation of photosynthetic acclimation and leaf N based on eco-evolutionary optimality. We have demonstrated that dynamic growth and a balanced plant C and N economy can be simulated under rapidly (seasonally) varying environmental conditions. This provides a way forward for informing widely used DGVMs and land vegetation components of Earth System Models with basic ecological principles and thus enabling reliable projections of the land carbon cycle in a future climate.

## Appendix A1 Vegetation growth phenology

Vegetation growth phenology is modelled as a state *f*_cold_ that is controlled by cold temperatures. A logistic function is used for representing an instantaneous suppression of growth. Once vegetation is exposed to low temperatures, a reduced growth (dormant) state is attained. This dormant state is modelled as a function of the minimum daily temperature *T*_min_ as

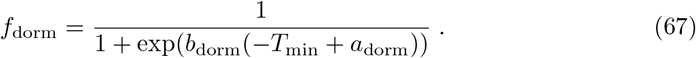

A new level of *f*_dorm_ is only attained if the new *f*_dorm_ is lower than the current *f*_dorm_, or if accumulated “warmth” (measured by growing degree days) triggers a release from the dormant state. The release is modelled as a function of growing degree days (GDD) as

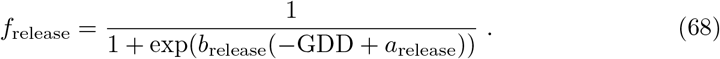

At each time step (day), the growth phenology state is updated, considering the entering into the dormant state and the release from it as

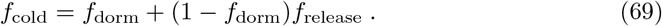

The summation of GDD is re-initiated each time vegetation is exposed to cold temperatures and *f*_cold_ is reduced. GDD is the sum of the difference between daytime mean temperature and a temperature threshold, here taken as 5^*°*^C. Parameters of Eqs. 67 and 68 are hard-coded (not treated as other calibratable model parameters (*a*_dorm_ = 2, *b*_dorm_ = 0.3, *a*_release_ = 150, *b*_release_ = 0.05).

**Figure A1.**
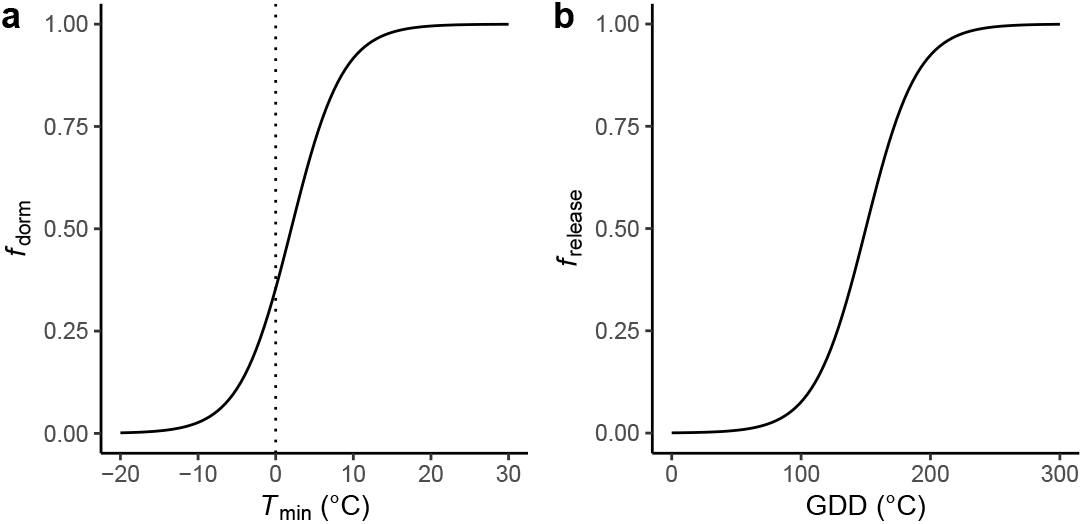
Vegetation growth phenology, represented by the response of the dormant state to daily minimum temperature (a) and the release from the dormant state as a function of growing degree days.

## Acknowledgments

BDS was funded by the Swiss National Science Foundation grant PCEFP2 181115. ICP acknowledges support from the ERC-funded project REALM (Re-inventing Ecosystem And Land-surface Models, Grant No. 787203). This work is a contribution to the LEMONTREE (Land Ecosystem Models based On New Theory, obseRvations and ExperimEnts) project, funded through the generosity of Eric and Wendy Schmidt by recommendation of the Schmidt Futures program (BDS, ICP).

## Code availability

The model is implemented as an R package in the {rsofun} modelling framework and is currently as branch cnmodel, publicly available at https://github.com/stineb/rsofun/tree/cnmodel.

